# IGFBP2 protects against pulmonary fibrosis through inhibiting P21-mediated senescence

**DOI:** 10.1101/2021.01.21.427684

**Authors:** Chin Chiahsuan, John Lee, Ranjith Ravichandran, Timothy Fleming, Stephen Wheatcroft, Mark Kearney, Ross Bremner, Thalachallour Mohanakumar, David J Flint, Angara Sureshbabu

## Abstract

Accumulation of senescent cells contributes to age related diseases including idiopathic pulmonary fibrosis (IPF). Insulin-like growth factor binding proteins (IGFBPs) are evolutionarily conserved proteins that play a vital role in many biological processes. Overall, little is known about the functions of IGFBP2 in the epigenetic regulation of cellular senescence and pulmonary fibrosis. Here, we show that *Igfbp2* expression was significantly downregulated at both mRNA and protein levels in a low-dose bleomycin-induced pulmonary fibrosis model of aged mice. Using the reduced representation of bisulfite sequencing technique, we demonstrated *Igfbp2* downregulation is attributed to DNA methylation of CpG islands in fibrotic lungs of aged mice. Furthermore, *Igfbp2* siRNA knockdown increased both P53 and P21 protein levels in mouse lung epithelial cells exposed to hypoxia treatment. Lentiviral mediated expression of *Igfb2* decreased P21 protein levels and significantly reduced beta galactosidase activity in mouse lung epithelial cells challenged with a senescent drug (atazanavir) and hypoxia treatments. Using the RT2 Profiler PCR Array, we found that P21, PAI-1, IRF-5 and IRF-7, important regulators of senescence pathway, were significantly downregulated specifically in type-II alveolar epithelial cells (AECs) of aged human-*Igfbp2* transgenic mice after bleomycin challenge. Finally, transgenic expression of human-*Igfbp2* in type-II AECs from aged bleomycin challenged mice significantly decreased senescent associated secretory phenotype factors and also reduced extracellular matrix markers compared to aged wild-type mice challenged with bleomycin injury. Collectively, these findings reveal that epigenetic repression of *Igfbp2* promotes pulmonary fibrosis and that restoring IGFBP2 in fibrotic lungs could prove effective in IPF treatment.

## Introduction

Idiopathic pulmonary fibrosis (IPF) is the most common and severe form of interstitial lung disease affecting more than 5 million individuals worldwide (1). IPF is a heterogeneous disease often occurring in the elderly and there are no effective treatments that can reverse or halt the disease. IPF has a dismal prognosis, with a median survival rate of less than 3 years. Advanced age is a significant risk factor for IPF and yet very little is known about how age contributes to the pathogenesis of IPF (2). Although much of the pathogenesis of IPF remains unclear, alveolar epithelium emerged as a principal site of injury in IPF. This disease is typically characterized with aberrant alveolar epithelium that triggers myofibroblast activation and excessive extracellular matrix deposition (3).

Cellular senescence has recently been implicated as a critical underpinning of age-related fibroproliferative diseases including IPF (4). Cellular senescence, a stress response associated with stable cell cycle arrest is accompanied by the robust secretion of proinflammatory cytokines leading to senescence-associated secretory phenotype (SASP) (5). Growing evidence suggests that P53 and P21 are two key molecular regulators in senescence pathway (6, 7) and, accumulation of P21 is typically observed during the process of senescence development (8). Recent studies have demonstrated that selective removal of senescent cells ameliorated a number of age-related diseases (4, 9, 10). However, the underlying mechanisms of cell senescence in alveolar epithelium remains poorly understood.

The insulin-like growth factor (IGF) system consists of 6 IGF-binding proteins (IGFBPs) with high affinity binding to IGF-I and IGF-II but not insulin (11-13). IGFBPs are ubiquitously expressed in most tissues and their actions are complex and depend upon the tissue and their ability to interact with proteins in the extracellular matrix (14). Although the actions of IGFs are modulated by IGFBPs, the cellular functions can also be affected via IGF-independent mechanisms (15, 16). IGFBP2 in particular may play an important role in lung function as IGFBP2 levels predicted the risk of a rapid evolution of systemic sclerosis induced interstitial lung disease (17). Growing evidence suggests that IGFBP2 undergoes epigenetic alterations in a tissue specific manner and predicts the development of chronic diseases.(18, 19) Further, IGFBPs play an important role in senescence and aging (20-22). So, we directed our focus on the role of IGFBP2 in epigenetic regulation of cellular senescence specifically in type-II alveolar epithelial cells (AECs) during the pathogenesis of idiopathic pulmonary fibrosis.

In the current study, we investigated whether IGFBP2 regulates cellular senescence and secretion of profibrotic mediators in type-II AECs in the context of idiopathic pulmonary fibrosis. We demonstrated that *Igfbp2* expression at both mRNA and protein levels significantly decreased in the lung homogenates of aged mice after low-dose bleomycin challenge. Utilizing gene silencing and overexpression approaches, we show that IGFBP2 inhibits P21 in murine lung alveolar epithelial cells upon hypoxia treatment. Utilizing bleomycin-induced aged pulmonary fibrosis model, we demonstrated that IGFBP2 regulates senescence markers in the type-II AECs of aged IGFBP2 transgenic mice. Finally, we report that transgenic expression of human-*Igfbp2* specifically in type-II AECs significantly reduced profibrotic mediators and extracellular matrix markers in response to low-dose bleomycin-induced pulmonary fibrosis in aged mice. These findings may provide new insights for the development of IGFBP-2 targeted therapy that could prove effective for IPF patients.

## Materials and Methods

### Animals

Aged 18-month-old (78 weeks) C57BL/6J mice were obtained from the Jackson Laboratories (Bar Harbor, Maine, USA). Human-*Igfbp2* tamoxifen inducible transgenic mice were obtained from Drs. Kearney and Wheatcroft (Leeds University, United Kingdom). A corresponding *Sftpc*-cre mouse line was obtained from Jackson Laboratories (Stock # 028054). Animals were housed under pathogen-free conditions with food and water ad libitum. All experiments and procedures were approved by the Institutional Animal Care and Use Committee at St. Joseph’s Hospital and Medical Center (Phoenix, Arizona).

### Bleomycin-induced pulmonary fibrosis

Aged (78 weeks or older) male and female C57BL/6J mice received intratracheal bleomycin (1 U/kg body weight) (Catalog # 203401; EMD Millipore, Burlington, Massachusetts, USA) or normal saline as previously described (23). To study the molecular signaling of IGFBP2, aged (26 weeks or older) human-*Igfbp2* transgenic mice and corresponding wild-type mice received intratracheal instillation of bleomycin (0.75 U/kg body weight) or saline. After the instillation, the anesthetized mice were kept on a warm bed for recovery. The experimental animals were monitored daily for adverse clinical signs, including body weight, appearance, hydration status, and any behavioral changes.

### Cell culture

MLE-12 (mouse lung epithelial) cell line was obtained from A.T.C.C. (American Type Culture Collection, Rockville, Maryland, USA), and cultured in DMEM/F12 medium enriched with 2% fetal bovine serum and 50 µg/ml plasmocin in a humidified atmosphere with 5% CO2 at 37 °C.

### Lentivirus transduction

Briefly, 1×10^4^ cells were mixed with mock or *Igfbp2* lentivirus (catalog # MR204287L3V; OriGene Technologies) at MOI=80 in the 500 ml of 10 µg/ml polybrene/DMEM-F12. The mixture was plated on the 24-well plate and centrifuged at 900g for 2 hours at 25°C. After spinfection, the extra 500 μl of 2%FBS/DMEM-F12 were added dropwise onto the cells. The plate was incubated for 48 hours at 37°C, and replaced the medium to 2% FBS/DMEM-F12. 0.5 µg/ml puromycin was used for the selection. The cells were lysed and the IGFBP2 level was estimated by immunoblotting.

### Immunoblotting

Total mice lungs or MLE-12 Cells were lysed by RIPA Lysis Extraction Buffer (catalog #89901; Thermo Fisher Scientific) along with protease and phosphatase inhibitors cocktail (catalog #78445; Thermo Fisher Scientific). Total protein concentration was determined by Pierce BCA protein assay reagent kit (catalog #23227; Thermo Fisher Scientific) according to the manufacturer’s protocol. 40 µg of total protein was loaded and separated by SDS-polyacrylamide gel electrophoresis. The gel was transferred using a Mini Trans-Blot cell (Biorad) to PVDF membrane (catalog #IPVH00010; EMD Millipore). Proteins were detected by mouse anti-IGFBP2 (R&D Systems, catalog #MAB7971), anti-P21 (Abcam, catalog #ab188224), anti-β-actin (catalog #sc-47778HRP; Santa Cruz Biotechnology) and anti-histone H3 (catalog #4499; Cell Signaling Technology). Immunoblots were incubated with SuperSignal™ West Pico or Femto Maximum Sensitivity Substrate (catalog #34095; Thermo Fisher Scientific).

### Statistical Analysis

All statistical tests were analyzed with the software (GraphPad Prism version 6.0). Statistical analysis was performed using 2-way ANOVA or 1-way ANOVA followed by Tukey’s post-hoc test. Student’s unpaired t-test was used to compare two groups. *P* value of less than 0.05 was considered statistically significant.

## Acknowledgements

We would like to thank Kristine Nally for proofreading the manuscript.

## Financial Statement

This study did not receive any funding from public or commercial or not-for-profit sources.

## Results

### Low-dose bleomycin induces non-resolving pulmonary fibrosis in aged mice model

Intratracheal administration of bleomycin is the most widely used model for studying pulmonary fibrosis in mice. However, several studies have utilized young mice in an attempt to recapitulate features of idiopathic pulmonary fibrosis (IPF). Since IPF is an age-related disease, we administered low-dose intratracheal bleomycin (1U/kg body weight) in an aged (78 weeks old) mice after intubation. Body weight was measured every week and found to be significantly decreased in bleomycin treated compared to normal saline at 7, 14 and 21 days (Fig. 1A). To detect fibrosis, lungs were evaluated for histological evidence; both Sirius Red and Trichrome staining increased after 28 and 50 days, indicating non-resolving fibrosis (Fig. 1B – 1C). The protein levels of collagen-I, fibronectin and vimentin were increased in the lung tissues of aged mice subjected to low-dose intratracheal bleomycin administration. Furthermore, bleomycin treatment decreased *Igfbp2* expression at both mRNA and protein levels but increased P21 protein levels (senescence marker), in concomitant with the activation of extracellular matrix (ECM) proteins (Fig. 1D – 1E).

**Figure 1.**
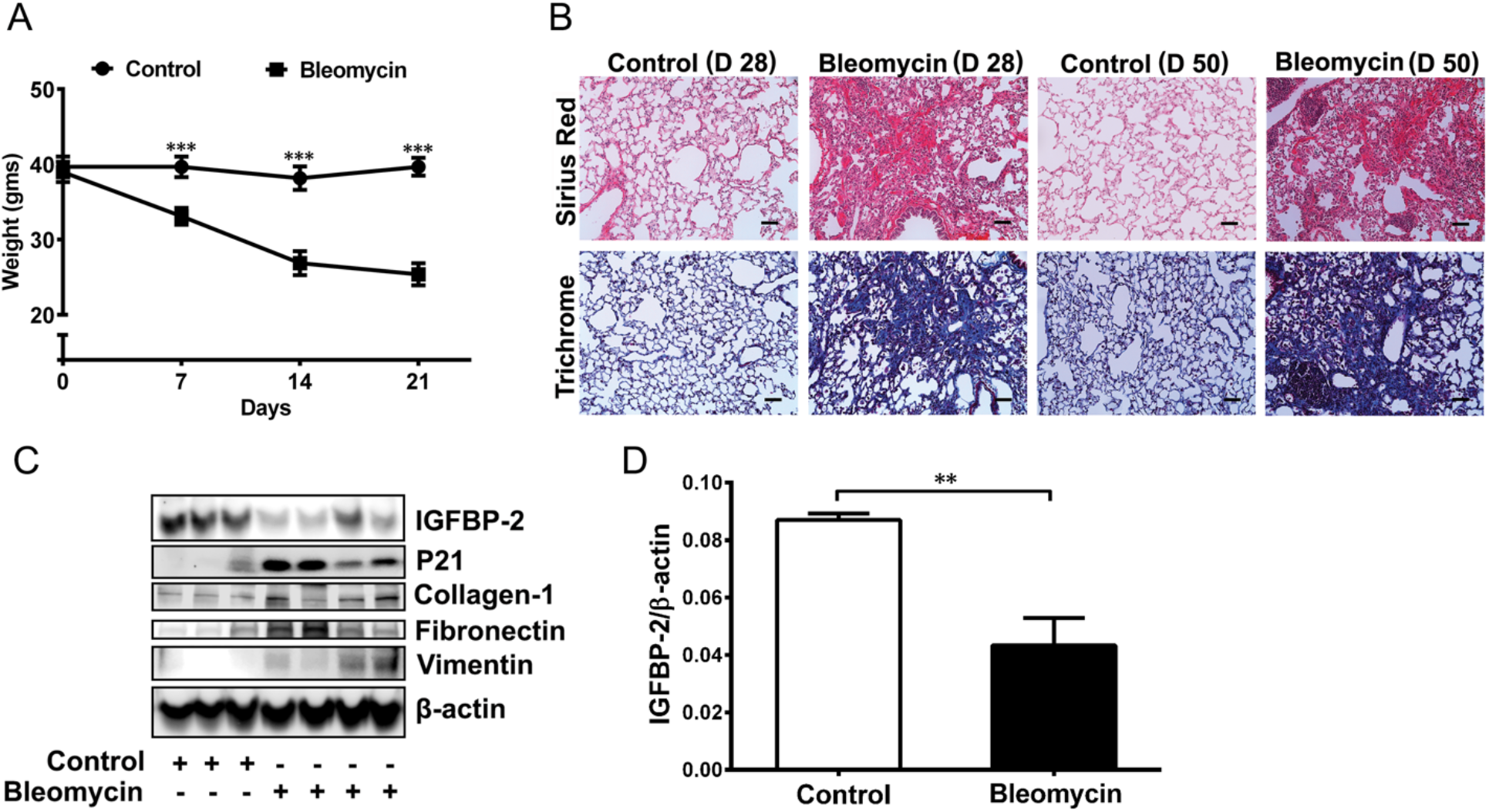
Low-dose bleomycin induces irreversible pulmonary fibrosis in aged mice. (A) Line plot showing body weights of aged (78 weeks old) wild-type (WT; C57BL/6J) mice 7, 14 and 21 days after intratracheal administration of normal saline (equal volume) or bleomycin (1U/kg body weight). (n = 8 WT saline; n = 8 WT bleomycin). (B-C) Sirius Red or Mason’s Trichrome stained lung sections of aged mice 28 or 50 days after intratracheal administration of bleomycin. Scale bars: 50 *μ*m. (n = 6 WT saline; n = 8 WT bleomycin) (D) Western blot for expression of IGFBP2, P21, Collagen-I, Fibronectin and Vimentin in the lung homogenates of aged mice 14 days after low-dose bleomycin challenge. Beta-actin is used as the internal control. (n = 6 WT saline; n = 8 WT bleomycin). (E) IGFBP2 mRNA expression in the lung homogenates of aged mice subjected to low-dose bleomycin challenge after 14 days. (n = 6 WT saline; n = 6 WT bleomycin). Data are mean ± SEM of 3 independent experiments. ***P < 0.001, two-way ANOVA **P < 0.01, Student’s unpaired t-test.

### Reduced representation of bisulfite sequencing profiling in aging model of pulmonary fibrosis

With the findings of decreased *Igfbp2* expression at both mRNA and protein levels, regulation is thought to occur at the DNA level. So, we performed global reduced representation bisulfite sequencing in the lung homogenates of aging mice challenged with bleomycin treatment. Bisulfite sequencing analysis revealed *Fnip2, Rnux1, Snx21, Hoxa3* and *Bc1* as some of the top differentially methylated genes in the aging model of bleomycin-induced pulmonary fibrosis (Fig. 2A – 2B). Interestingly, we also observed significantly increased DNA methylation of *Igfbp2* but no change in DNA methylation of *Cdkn1a* (also known as P21) in aged low-dose bleomycin treated lungs as compared to saline control suggesting a regulatory role for *Igfbp2* gene methylation (Fig. 2C – 2E).

**Figure 2.**
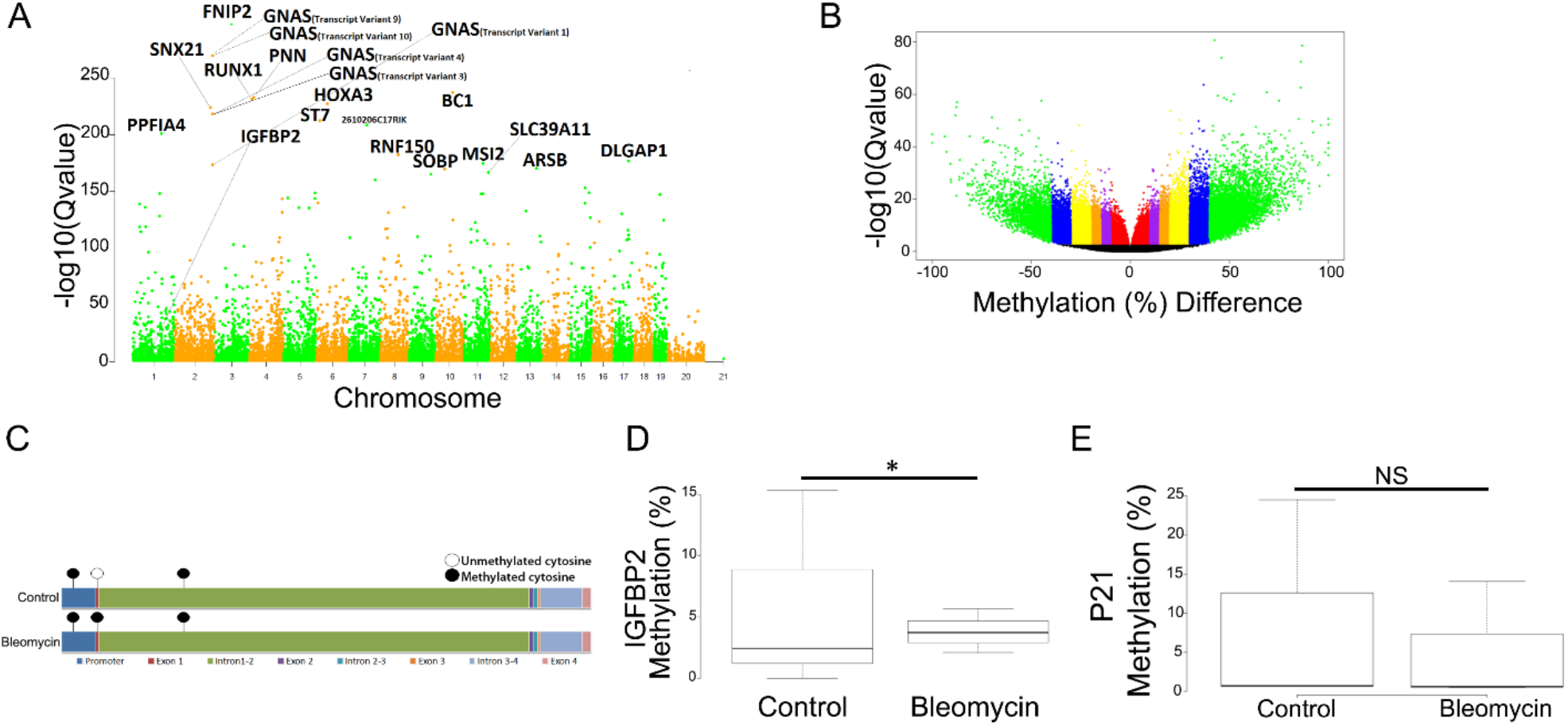
Reduced Representation of Bisulfite Sequencing of low-dose bleomycin induced pulmonary fibrosis in aged mice. (A) Manhattan plot showing the top 20 differentially methylated genes and IGFBP2 gene for all the chromosomes. The red line indicates the significant level of Q-value < 0.01. (B) Volcano plot showing the methylation (%) difference in all genes. (C) Lollipop plot showing the methylated cytosine regions of IGFBP2 promoter, exons and introns. (D) Box plot showing the significantly increased differentially methylation region of *Igfbp2* variant (NM_001310659.1) in low-dose bleomycin-induced pulmonary fibrosis mice. (E) Box plot showing no change of differentially methylated region of *Cdkn1a* in low-dose bleomycin-induced pulmonary fibrosis mice. (n = 3 WT saline; n = 3 WT bleomycin) *P < 0.05 Student’s unpaired t-test.

### IGFBP2 deficiency elevates senescent markers upon hypoxia treatment

To assess the biological relevance of IGFBP2, we utilized mouse lung epithelial (MLE-12) cells exposed to 0.1% hypoxia as an in vitro model of alveolar epithelial injury. The protein levels of P53 and P21 were increased but IGFBP2 protein levels decreased in MLE-12 cells exposed to 0.1% hypoxia at 4 h (Fig. 3A). Furthermore, we showed decreased IGFBP2 protein levels in the nuclear fractions of MLE-12 cells challenged with hypoxia as compared to normoxia (Fig. 3B). Furthermore, to evaluate the function of IGFBP2 we performed siRNA transfection in MLE-12 cells challenged with hypoxia treatment. We showed increased P53 and P21 protein levels in MLE-12 cells treated with *Igfbp2* siRNA compared to non-targeting siRNA and that increase was also observed after 4 h hypoxia treatment (Fig. 3C).

**Figure 3.**
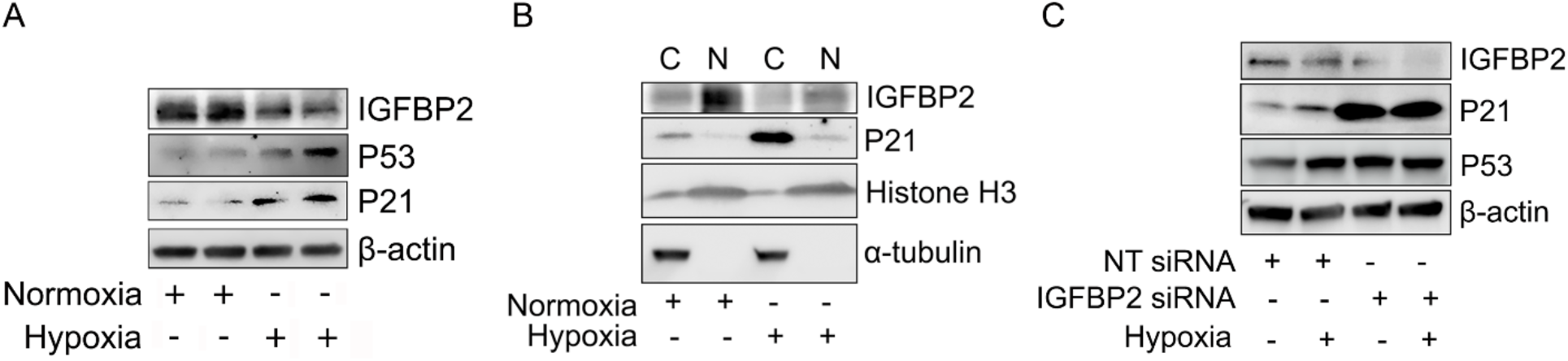
IGFBP2 deficiency increases P21 expression in response to hypoxia stimulus (A) Western blot for the expression of IGFBP2, P53, P21 in MLE-12 cells exposed to absence or presence of hypoxia treatment. (B) Western blot for the expression of IGFBP2 and P21 in the cytosolic and nuclear fractions of MLE-12 cells that were exposed to absence or presence of hypoxia treatment. Alpha-tubulin and histone-3 served as internal controls (C) Non-targeting or *Igfbp2* siRNA transduced MLE-12 cells were exposed to absence or presence of hypoxia treatment. Western blot for the expression of IGFBP2, P53 and P21. Beta-actin served as internal control. Data are representative of minimum of 3 independent experiments.

### Lentiviral expression of IGFBP2 reduces P21 expression and beta-galactosidase activity

To determine the action of IGFBP2, we undertook a lentiviral transduction approach in MLE-12 cells exposed to hypoxia treatment. Consistent with the results from *Igfbp2* silencing, lentiviral expression of *Igfbp2* decreased P21 levels in MLE-12 cells in response to hypoxia treatment when compared with mock virus treated MLE-12 cells exposed to hypoxia (Fig. 4A). Furthermore, both cytosolic and nuclear fractions showed decreased P21 levels in *Igfbp2* lentivirus-transduced MLE-12 cells in response to hypoxia treatment (Fig. 4B).

**Figure 4.**
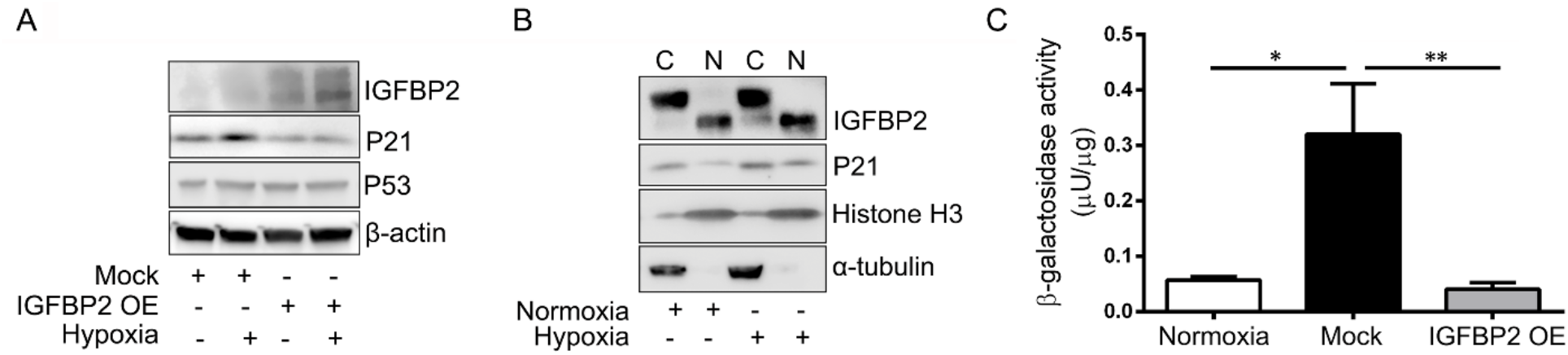
Stable transduction with *Igfbp2* lentivirus vector decreased P21 expression and beta-galactosidase activity (A) Mock virus and *Igfbp2* lentivirus-transduced MLE-12 cells in the absence or presence of hypoxia treatment. Western blot for the expression of IGFBP2, P21 and P53. Beta-actin served as internal control (B) Western blot for the expression of IGFBP2 and P21 in the cytosolic and nuclear fractions of *Igfbp2* lentivirus-transduced MLE-12 cells in the absence or presence of hypoxia treatment. (C) Bar graph showing the beta galactosidase activity of MLE-12 cells treated with Atazanavir and hypoxia for 96 h. Data are representative of minimum of 3 independent experiments.

Since senescence-associated (SA) β-galactosidase (*β*-gal) activity is a marker for senescence, we evaluated the endogenous SA-β-gal function of IGFBP2 in MLE-12 cells exposed to atazanavir (protease inhibitor) and hypoxia treatments. Biochemical analysis showed that SA-β-gal activity significantly increased in MLE-12 cells after 4 days of treatment with atazanavir and hypoxia. Importantly, *Igfbp2* expressed through lentivirus transduction significantly decreased SA-β-gal activity in MLE-12 cells after treatment with atazanavir and hypoxia at 96 h (Fig. 4C).

### Senescence RT2 Profiler PCR Array in type-II alveolar epithelial cells of human-IGFBP2 transgenic mice

To study the mechanism of IGFBP2 in the development of pulmonary fibrosis, we utilized tamoxifen inducible transgenic (Tg) mice that have the human-*Igfbp2* gene knocked into the ROSA26 locus flanked by a floxed STOP codon (Fig. E 1A – 1B). Utilizing RT2 Profiler PCR Array, first we examined the expression of senescence genes in the aged (78 weeks old) lungs of low-dose bleomycin treated wild-type mice. We identified significant upregulation of important senescence genes — *Cdkn2a, Cdkn2b, Ifn-γ, Pai-2, Tp53bp1* along with 34 other senescent related genes (Fig. E 2A – 2B). To investigate the effects of *Igfbp2*, primary type-II AECs were isolated from human-*Igfbp2* Tg mice after bleomycin injury. Mouse cellular senescence RT2 Profiler PCR Array was used to analyze the expression of senescence genes specifically in type-II AECs of aged (26 weeks old) human-*Igfbp2* Tg mice after 14 days of bleomycin treatment. qPCR array data analysis identified 31 upregulated and 21 downregulated genes relevant to cellular senescence pathway. Primary type-II AECs from aged human-*Igfbp2* Tg mice exposed to bleomycin treatment showed downregulation of important senescence genes viz., *Cdkn1a* (P21), *Pai-1, Irf-5, Irf-7, Tp53bp1* as well as fibronectin as compared to aged wild-type mice exposed to bleomycin treatment (Fig. 5).

**Figure 5.**
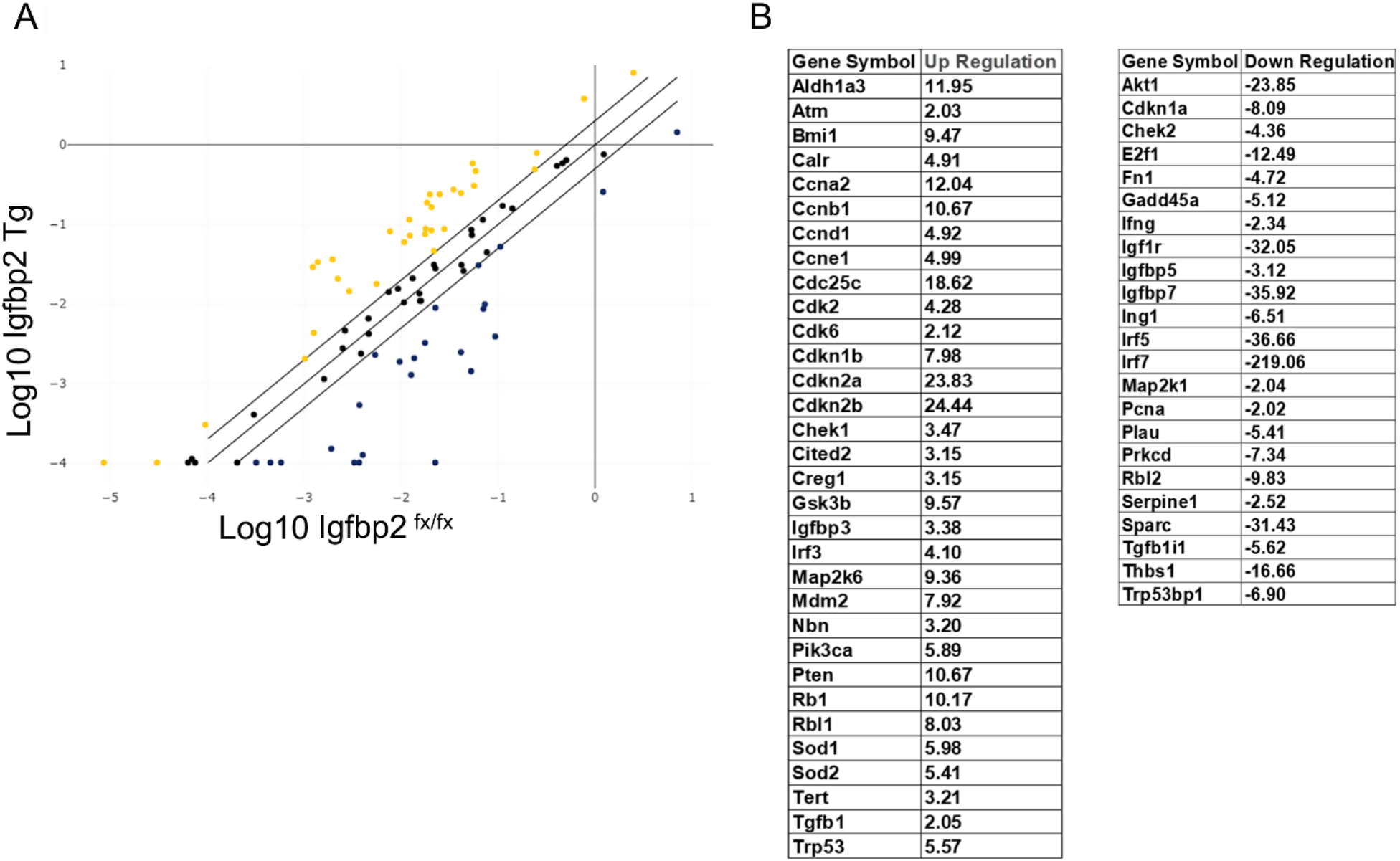
Cellular senescence gene profiling in type-II alveolar epithelial cells of aged human-*Igfbp2* transgenic mice (A) Scatter plot showing the upregulated and downregulated genes relevant to senescence pathway in isolated type-II AECs from aged (26 weeks old) human-*Igfbp2* transgenic mice after 14 days of low-dose bleomycin treatment (0.75U/kg body weight). Yellow dots indicate upregulated genes; black dots indicate no change; blue dots indicate downregulated genes. (B) Table showing the list of fold regulated genes with gene symbol. + indicates upregulated genes; -indicates downregulated genes. (n = 3 *Igfbp2* fx/fx; n = 3 *Igfbp2* Tg)

### Transgenic expression of human-IGFBP2 reduces extracellular matrix deposition and senescence associated secretory phenotype

We next examined the in vivo effect of human-*Igfbp2* transgenic aged mice, 6 months or older, in bleomycin-induced pulmonary fibrosis. To this end, we first assessed bleomycin-induced weight loss at 7 day intervals after intratracheal instillation of bleomycin injury. Human-*Igfbp2* transgenic mice were significantly protected from weight loss at days 7, 14 and 21 as relative to aged wild-type mice after bleomycin injury (Fig. 6A). Both Sirius Red and Trichrome staining showed reduced lung fibrosis content in human-*Igfbp2* transgenic mice at 28 days as relative to aged WT mice challenged with bleomycin treatment (Fig. 6B). Similarly, compared with aged wild-type mice, human-*Igfbp2* transgenic mice showed decreased extracellular matrix markers — Collagen-1, Fibronectin and Vimentin in the lungs at 14 days after bleomycin injury (Fig. 6C – 6D). Furthermore, quantitative RT-PCR analysis of type-II AECs from aged human-*Igfbp2* mice showed downregulation of SASP factors viz., *Il-1β, Tnf-α, Mcp-1, Stat6* and *Il-4* compared to aged wild-type mice challenged with bleomycin treatment indicating abrogation of pulmonary fibrosis (Fig. 6E).

**Figure 6.**
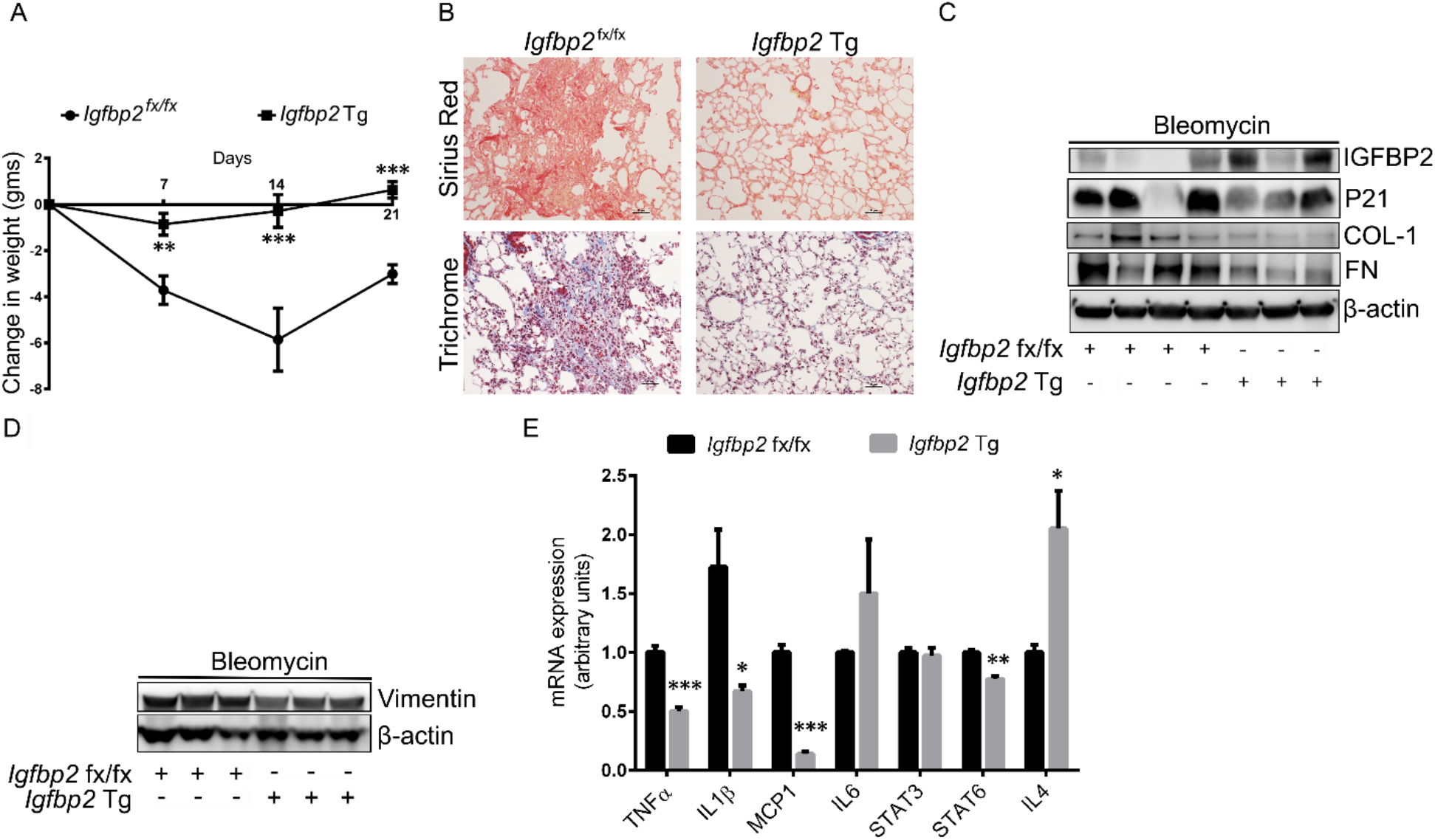
Effects of aged human-*Igfbp2* transgenic mice challenged with bleomycin treatment. (A) Line plot showing the change in body weights of aged (26 weeks) wild-type and human-*Igfbp2* transgenic mice subjected to intratracheal administration of bleomycin treatment (0.75U/kg bodyweight). (n = 7 *Igfbp2* fx/fx; n = 7 *Igfbp2* Tg). ***P < 0.001 and **P < 0.01, two-way ANOVA (B) Sirius Red or Mason’s Trichrome stained lung sections of aged wild-type and human-*Igfbp2* transgenic mice 28 days after intratracheal administration of bleomycin treatment. Scale bars: 50 *μ*m. (n = 4 *Igfbp2* fx/fx; n = 3 *Igfbp2* Tg). (C-D) Western blot for the expression of Collagen-I, Fibronectin, P21 and Vimentin. (n = 4 *Igfbp2* fx/fx; n = 3 *Igfbp2* Tg). (E) RT-PCR analysis for mRNA expression of TNF-*α*, IL-1*β*, MCP-1, IL-6, STAT3, STAT6 and IL-4 in aged wild-type and human-*Igfbp2* transgenic mice 14 days after intratracheal administration of bleomycin. (n = 4 *Igfbp2* fx/fx; n = 3 *Igfbp2* Tg). ***P < 0.001, **P < 0.01 and *P < 0.05 Student’s unpaired t-test.

## Discussion

In the present study we provide evidence that IGFBP2 inhibits P21, well established senescence marker, and reduces non-resolving pulmonary fibrosis in aged mice after bleomycin injury. Using type-II AEC specific *Sftpc* promoter human-*Igfbp2* transgenic mice, we are capable of demonstrating reduced senescence, SASP factors, extracellular matrix deposition and improved pulmonary fibrosis in the bleomycin-challenged aged mice.

Cellular senescence, an evolutionarily selected process, counteracts early life cancer (24). Initially, senescence may act as protective response by limiting proliferation but studies have shown that accumulation of senescence is the key pathogenic mechanism that drives the development of pulmonary fibrosis (4, 25). Advanced age is the predominant risk factor for age related pathologies and importantly for the development of pulmonary fibrosis (26). Recent studies have utilized aged mice as an improved new model for idiopathic pulmonary fibrosis (27). In line with this, we confirm that challenging aged mice (18 months old), with low-dose bleomycin injury resulted in non-resolving pulmonary fibrosis even after 50 days.

Studies have indicated that senescence is also accompanied by epigenetic and chromatin remodeling of nuclear architecture (28, 29). In this study, DNA methylation of *Igfbp2* was significantly increased in bleomycin-challenged aged mice but DNA methylation of *Cdkn1a* (P21) did not change. Consistent with this, recent reports demonstrated upregulation of P21 expression in human lungs with IPF compared to donor lungs (30, 31). In contrast, decreased expression of IGFBP2 specifically in fibrotic lungs of aged mice may seem counterintuitive to a recent report showing higher levels of circulating IGFBP1 and IGFBP2 in the serum of IPF patients (32). However, the elevated serum levels of IGFBP2 in IPF patients may be reasoned due to aberrant secretion by immune cells eliciting an immune response and is a subject of further investigation. Another possible explanation is that the relationship of *Igfbp2* DNA methylation in lungs and circulating IGFBP2 levels are largely independent i.e., heterogenous cell-types involved and functional roles of IGFBP2 acting downstream of IPF.

Alveolar epithelial cell senescence is evident in IPF and in a variety of experimental models of lung fibrosis (30). P53 plays a central role in cellular senescence mainly through P21 induction. In this study, we show that silencing *Igfbp2* in mouse lung epithelial cells increased both P53 and P21 protein levels in response to hypoxia treatment. Similarly, lentiviral mediated expression of *Igfbp2* decreased P21 protein levels and also reduced SA-*β*-galactosidase activity in response to atazanavir and hypoxia treatments. There is considerable evidence that IGFBP2 is detected in the nuclei and such nuclear translocation is mediated by a functional nuclear localization signal sequence (33). In relevance to this, we show that IGFBP2 protein levels were upregulated in the nuclear fraction as relative to the cytosolic fraction in MLE-12 cells exposed to hypoxia treatment.

Previous studies have shown that type-II AECs undergo senescence and secrete profibrotic mediators in bleomycin-induced pulmonary fibrosis (34). In this study, we systematically examined senescence markers in type-II AECs obtained from human-*Igfbp2* transgenic mice. We show downregulation of senescent markers — *Cdkn1a*, encoding P21, *Pai-1, Irf-5, Irf-7* and *Tp53bp1* in aged human-*Igfbp2* transgenic mice as compared to aged wild-type mice challenged with bleomycin treatment. In agreement with this, a recent report showed interferon regulated genes — IRF-5 and IRF-7 induce senescence in immortal fibroblasts (35). Another report showed increased P53BP1 positive foci in the development of radiation-induced lung injury (36). Importantly, emerging evidence suggests that DNA damage-induced P21 expression increasing to stable levels allows the cell to permanently exit the cell cycle and undergo senescence or apoptosis (6, 37). In addition to senescence related genes, we investigated the expression of SASP in type-II AECs overexpressing human-*Igfbp2* after bleomycin-induced lung injury. Specifically, we show significant downregulation of established SASP components — *Tnf-α, Il-1β, Mcp-1* and *Stat6*. Although *Il-4* was upregulated, recent reports suggest that IL-4 plays a dichotomous role in pulmonary fibrosis (38, 39). Nevertheless, we demonstrated that IGFBP2 regulates both P21 levels and profibrotic mediators in bleomycin-challenged aged mice suggesting that IGFBP2 acts as an upstream regulator of P21-mediated senescence. Furthermore, we show reduced protein levels of ECM markers in the lungs of human-*Igfbp2* transgenic mice compared to aged WT mice challenged with bleomycin exposure. These data support the concept that restoring IGFBP2 levels in the fibrotic lungs may have beneficial effect for IPF patients.

In summary, we showed that IGFBP2 inhibits P21-mediated senescence at least in type-II AECs and also reduces SASP components in aged mice subjected to bleomycin-induced lung injury and fibrosis. These results suggest that transgenic expression of human-*Igfbp2* specifically in type-II AECs reduces non-resolving pulmonary fibrosis by inhibiting senescence as well as profibrotic mediators.

## Online Data Supplement

## Methods

### Hypoxia treatment

Approximately 16 h before the exposure to hypoxia, 1×10^6^ MLE-12 were seeded on to the 10 cm dish. Subsequently, MLE-12 cells were transferred to the incubation chamber with 0.1% O_2_ and 5% of CO_2_ for the different time points exposure. After hypoxia treatment, cells were prepared for downstream experiments.

### Generation of SFTPC-cre-ERT2-human-Igfbp2 transgenic mice

Sftpc (surfactant protein C)-cre hIgfbp2^flox^ (human Igfbp2 knocked into the ROSA26 locus flanked by a floxed STOP codon) was generated by crossing a mouse line expressing tamoxifen-inducible cre under control of the type II alveolar epithelial cell-specific promoter Sftpc with human-Igfbp2^flox^. The genotype of transgenic mice was confirmed by PCR with the following primers:

182053 Cre1: 5′-CTCCCAAAGTCGCTCTGAGTTGTTATCA – 3′

182054 Cre2: 5′-CGATTTGTGGTGTATGTAACTAATCTGTCTGG-3′ 0034-Kin-ROSA-GX6044:5′-GCAGTGAGAAGAGTACCACCATGAGTCC-3′

13007-mutant reverse: 5′-ACACCGGCCTTATTCCAAG-3′

24999-common: 5′-TGCTTCACAGGGTCGGTAG-3′

25000-Wild type reverse: 5′-TGCTTCACAGGGTCGGTAG-3′

The band was at 245 bp for the homozygous of Igfbp2, and 210 bp for the Sftpc-cre driven mice. To induce the Cre expression, mice were injected via i.p. with 5 doses of tamoxifen every other day for 0.45 mg/kg body weight.

### Reduced representation of bisulfite sequencing analysis

Total lung DNA from aged (78 weeks old) wild-type mice challenged with bleomycin (1U/kg body weight) treatment were extracted using DNeasy Blood & Tissue kit (catalog # 69504, Qiagen). DNA concentration was assessed using Nanodrop and only samples with optical density A260/A280 <u>≥</u>1.8 were chosen. Reduced Representation of Bisulfite Sequencing (RRBS) was performed on a RRBS service basis by the company Diagenode (Diagenode Inc, catalog # G02020000). The comparisons between the saline (control) and bleomycin treatment RRBS datasets were carried out using R package GeneDMR (1). RefGene database mm10 and CpG island annotation from the UCSC genome browser (http://genome.ucsc.edu) were used. Differentially methylated regions (DMRs) were annotated using the UCSC Refseq tracks (mm10) to further analyze CpG sites included in genes — promoter regions, introns and exons upstream and downstream of transcriptional start site.

### Cytosolic/ nuclear protein fractionation

1.0×10^7^ MLE-12 cells in the 10 cm dish were exposed to the hypoxia condition for 4 and 24 hours, respectively. Cells were harvested, and resuspended with 1x cytosolic extraction buffer (10 mM HEPES, 1.5 mM MgCl2, 10 mM KCl, 0.05% NP40 pH 7.9). Then, cells were incubated on ice for 10 mins, and lysed with 25 µl Thermo Scientific™ RIPA Lysis and Extraction Buffer (Thermo Fisher Scientific, catalog #89901) with protease and phosphatase inhibitor cocktails (Fisher Scientific, catalog #78445). The supernatant was collected after centrifugation for 10 mins at 800xg. The pellet was washed twice with 1x cytosolic extraction buffer. The pellet was lysed with 50 µl Thermo Scientific™ RIPA Lysis and Extraction Buffer and incubated on ice for 30 minutes by vortexing for 5-minute intervals. The nuclear fraction was collected from the supernatant after the centrifugation for 30 minutes at 14,000xg. The cytosolic/nuclear fraction was detected by immunoblotting assay.

### β-galactosidase activity assay

MLE-12 cells were seeded at 8×10^5^ cells per 10 cm dish at 37°C for overnight, and 20µM atazanavir (Sigma-Aldrich, catalog #SML1796-5MG) was added two times at 24 h intervals. Simultaneously, MLE-12 cells were incubated in hypoxia (0.1%) for 96 h. Endogenous β-galactosidase activity was measured by the β-Gal Activity Assay Kit (BioVision, catalog# K821-100). Briefly, MLE-12 cells were lysed using 100 μl ice cold β-Gal assay buffer for 15 min after atazanavir and hypoxia treatments for 96 h. Supernatant was collected after centrifugation at 10,000 X g for 10 min at 4°C. About 10 µl of supernatant was used along with β-Gal substrate and fluorescence was measured using Spectramax i3 fluorometer (Molecular Devices, Inc.) in kinetic mode for 5 – 60 min at 37 °C.

### Primary type II alveolar epithelial cell isolation

The Sftpc-Cre-ERT2-human-IGFBP2^flox^ transgenic mice and corresponding wild-type mice were sacrificed after 14 days of bleomycin (0.75 U/kg body weight) treatment, and the lung was perfused with 30 ml of saline to remove the blood. Lungs collected from the same condition mice were minced into 1 mm pieces and digested with the mixture of 1 mg/ml of collagenase I and 5 U/ml of Dispase at 37°C for 25 minutes. Cells were passed through a 400 µm filter and neutralized by equal volume of 20% FBS. Cells were centrifuged at 180xg and resuspended by 10% FBS with DMEM/F12. First, suspended cells were separated by CD45 MACS cell separation magnetic beads (catalog # 130-052-301, Miltenyi Biotec), and CD45 depletion cells were collected. CD45-cells were sequentially separated by EpCAM magnetic beads (catalog #130-105-958, Miltenyi Biotec). The column was washed three times with 3 ml of PBS containing 0.5% Bovine serum albumin (catalog #A30075, RPI Research), and EpCAM positive cells were collected after removing the column from the magnetic separator (catalog #130-091-051, Miltenyi Biotec) and flushing the column with 0.5% bovine serum albumin in the saline. CD45-/EpCAM+ cells were further filtered with a 30 µm filter and the total RNA was extracted from the cells by all prep DNA/RNA mini kit (catalog #80204, Qiagen).

### Senescence gene expression profiling

Cellular senescence gene expression profiling was performed using RT^2^ Profiler PCR Array (catalog #330231, Qiagen). AEC2 cells were isolated from the lungs of normal saline or bleomycin treated mice. RNA was extracted as per the manufacturer instructions described in RT^2^ first strand kit (catalog #330401, Qiagen). cDNA was combined with RT^2^ SYBR Green qPCR Master Mix (catalog #330520, Qiagen), and ΔΔCT was measured by StepOnePlus™ Real-Time PCR System (catalog #4376600, Thermo Fisher Scientific). The further analysis and fold regulation were calculated based on the ΔΔCT method using the GeneGlobe data analysis web portal (www.qiagen.com/geneglobe). The primers for senescence associated secretory phenotype (SASP) genes were obtained from RealTime Primers (# OS1, realtime primers.com). The list of primer sequences for the quantitative PCR for the SASP genes are below:

**Table E1:**
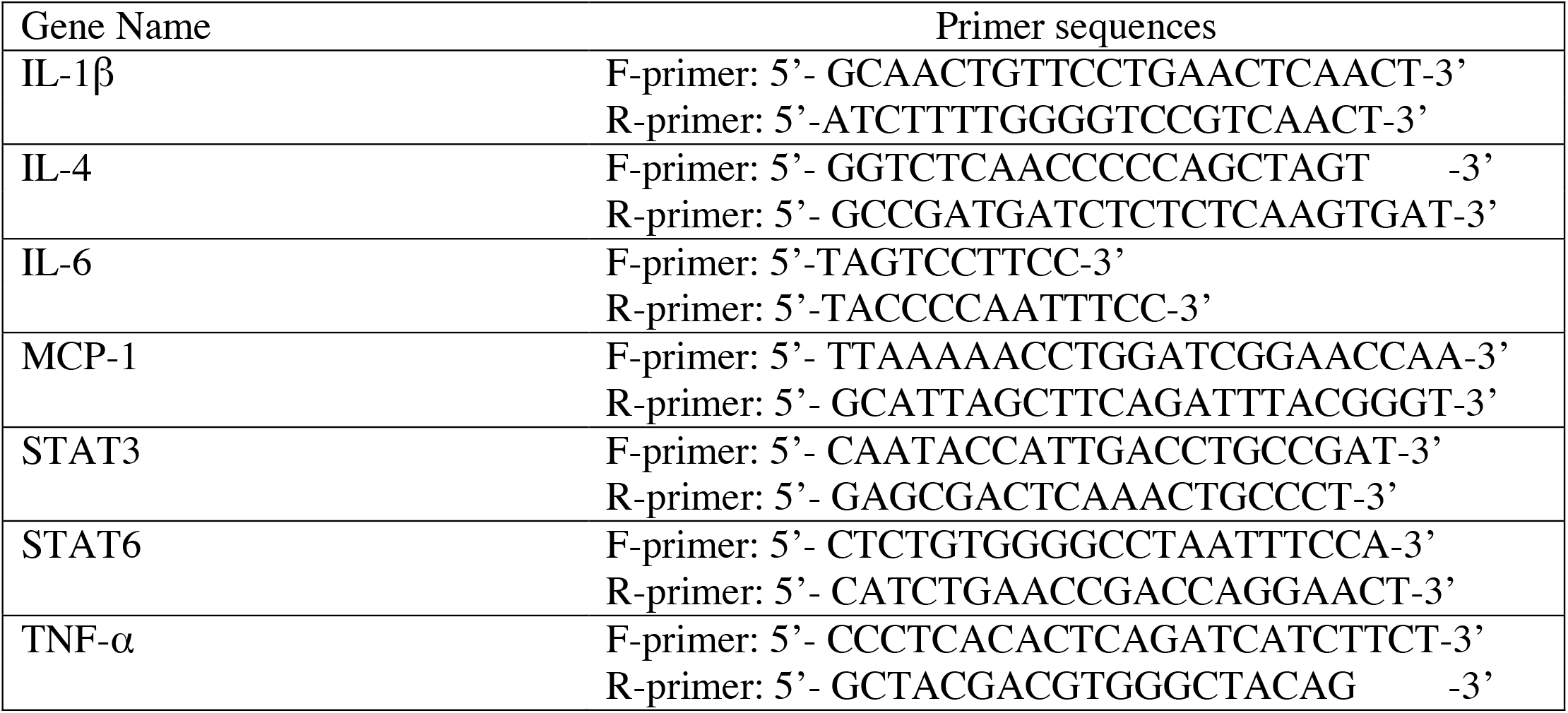
Mouse primer sequences for Qpcr.

## Supplementary Figures

**Figure E1.**
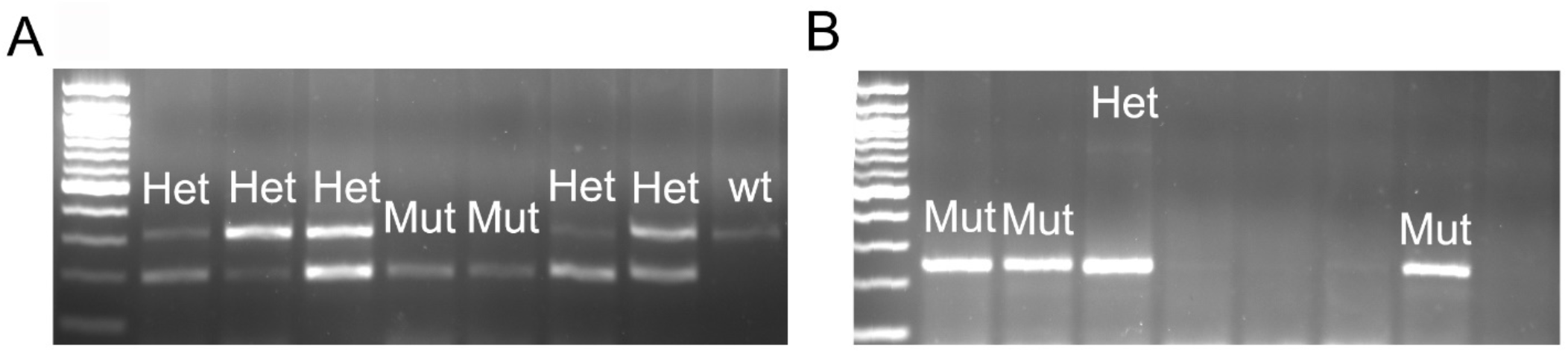
Agarose gel electrophoresis showing the genotyping of Igfbp2 transgenic mice obtained through crossing the Igfbp2 flox with Sftpc-cre mice. (A) Sequence of the PCR amplified regions encompassing wild-type, heterozygote and mutant genotypes for Sftpc-cre. Mutant, 327 bp; Heterozygous, 210 and 327 bp; wild-type, 327 bp (B) Sequence of the PCR amplified regions encompassing wild-type, heterozygote and mutant genotypes for Igfbp2 flox. Mutant, 245 bp; Heterozygous, 245 and 778 bp; wild-type, 778 bp.

**Figure E2.**
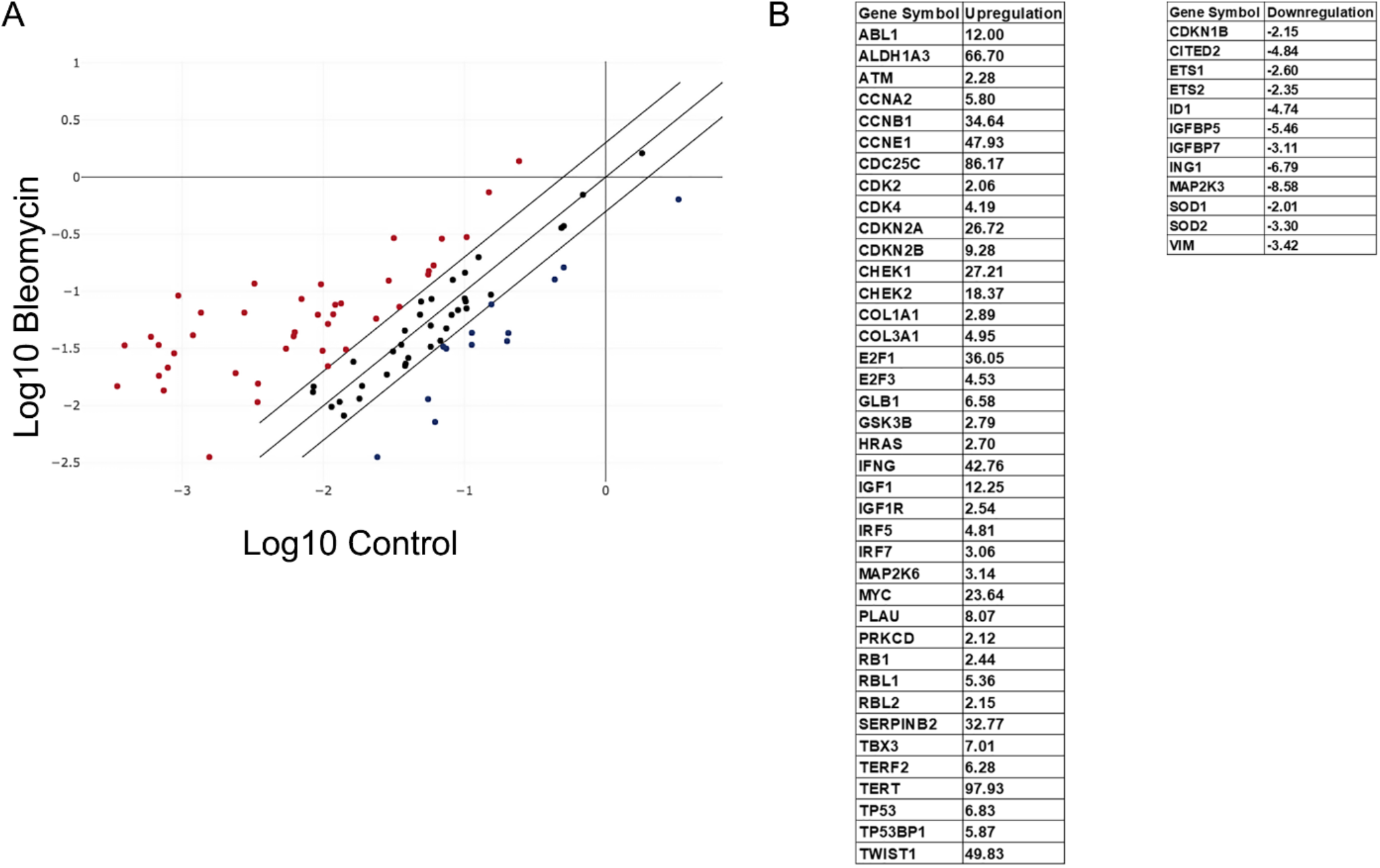
(A) Scatter plot showing the RT2 Profiler PCR Array for 84 genes related to cellular senescence pathways performed in the lungs of aged wild-type mice subjected to low-dose bleomycin injury. Red dots indicates upregulated genes; black dots indicate no change; blue dots indicate downregulated genes. (B) Table showing the list of fold upregulated and down regulated genes in the total lungs challenged with bleomycin as compared to normal saline (n = 3 saline control; n = 3 bleomycin).

